# Expanding the Coverage of the Metabolic Landscape in Cultivated Rice with Integrated Computational Approaches

**DOI:** 10.1101/2020.03.04.976266

**Authors:** Xuetong Li, Hongxia Zhou, Ning Xiao, Xueting Wu, Yuanhong Shan, Longxian Chen, Cuiting Wang, Zixuan Wang, Jirong Huang, Aihong Li, Xuan Li

## Abstract

Genome-scale metabolomics analysis is increasingly used for pathway and function discovery in post-genomics era. The great potential offered by developed mass spectrometry (MS)-based technology has been hindered by the obstacle that only a small portion of detected metabolites were identifiable so far. To address the critical issue of low identification coverage in metabolomics, we adopted a deep metabolomics analysis strategy by integrating advanced algorithms and expanded reference databases. The experimental reference spectra, and *in silico* reference spectra were adopted to facilitate the structural annotation. To further characterize the structure of metabolites, two approaches, structural motif search combined with neutral loss scanning, and metabolite association network were incorporated into our strategy. An untargeted metabolomics analysis was performed on 150 rice cultivars using Ultra Performance Liquid Chromatography (UPLC)-Quadrupole (Q)-Orbitrap mass spectrometer. 1939 of 4491 metabolite features in MS/MS spectral tag (MS2T) library were annotated, representing an extension of annotation coverage by an order of magnitude on rice. The differential accumulation patterns of flavonoids between *indica* and *japonica* cultivars were revealed, especially O-sulfated flavonoids. A series of closely-related flavonolignans were characterized, adding further evidence for the crucial role of tricin-oligolignols in lignification. Our study provides a great template in the exploration of phytochemical diversity for more plant species.

## Introduction

It is estimated that there are from 200,000 to 1,000,000 metabolites produced in green plants, underlying their broad chemical diversity and metabolic complexity [1]. Genome-scale metabolomics analysis has become a powerful tool in the elucidation of functional gene and pathway for diverse phytochemicals [2-5]. The more recent progresses in UPLC coupled with high-resolution MS, allow detecting metabolites at unparalleled sensitivity, resolution, accuracy, and throughput [6]. However, the great power in advanced liquid-phase separation and mass spectrometry technology has been limited, considering a vast majority of metabolite features detected from plants remain unidentified in current status [7, 8]. It is a major challenge to detect and identify the massive amount of heterogeneous phytochemicals with high dynamic range in concentrations, chemical and physical properties, and structures. The lagging in identification of metabolites from plant sources can be attributed to various factors, *e.g.*, the insufficient performance of early MS-based platforms, the structural complexity of diverse metabolites, the limited availability of reference mass spectra from standard compounds, and the low throughput for processing and structure elucidating of mass spectral data [9-12]. It is critical to handle and resolve the metabolomics data efficiently, in order to bridge the gap between technological advance and demands of plant metabolomics research. In recent years, progresses have been made in the improvement of metabolite annotation coverage through collecting reference mass spectra from more standard compounds [13-16], and developing computer-assisted approaches to facilitate the structure elucidation of metabolites [17-20].

Rice (*Oryza sativa* L.) is one of the major staple foods worldwide, and it is critical to explore its chemical compositions and metabolic traits for the enhancement of grains quality and nutritional value [21, 22]. The two major subspecies of cultivated rice, *indica* and *japonica*, formed during domestication, display distinct features in morphology and physiology [23-25]. In recent years, a series of studies on rice metabolomics were performed, which provides the foundation for the metabolic components of rice [2, 5, 26, 27]. However, there are plenty of unknown metabolite features in above studies and the metabolic diversity of rice still needs further efforts to explore. Other studies focused on phytochemical genomics to dissect the underlying genetics basis of biosynthesis and physiological function of metabolites during the evolution and adaptation of plants [28]. The metabolic quantitative trait loci (mQTL) mapping and metabolic genome-wide association study (mGWAS) were used to reveal the genetic polymorphisms and candidate genes that affected metabolic traits in rice [2, 5, 27, 29].

Our current study was designed to address the key issue in plant metabolomics, that is, the low identification coverage of metabolites. We sought to expand the annotation coverage with computational approaches, by adopting a deep metabolomics analysis strategy that combines experimental and *in silico* reference mass spectral libraries, and advanced algorithms. The structural motif search combined with neutral loss scanning and metabolite association network methods were integrated in our strategy to facilitate the characterization of structure and potential function of novel metabolites without reference from above libraries. As a proof-of-concept study, using state-of-the-art UPLC-Q-Orbitrap mass spectrometer platform, we performed an untargeted metabolomics analysis on a core collection of 150 *indica* and *japonica* cultivars grown in northeastern and southeastern China. A MS2T library for rice grains was constructed containing 4491 metabolite features, and of which 1939 were annotated. The annotation coverage of rice metabolome was significantly improved through our strategy. Further, our analyses revealed the systematic difference of metabolomes between *indica* and *japonica* subspecies and major differential accumulation patterns of flavonoid derivatives, especially O-sulfated flavonoids. A group of closely-related flavonolignans were newly uncover in rice, which provided further evidence for the crucial role of tricin-oligolignols in lignification of monocots. Our deep metabolomics analysis strategy expanded our understanding of phytochemical diversity and function in rice, which has profound implication for improving the quality and nutritional value of crops through genetic breeding.

## Results and discussion

### Integrated computational approaches and their evaluations

To handle the mass spectral data generated from UPLC-Q-Orbitrap mass spectrometer, we adopted a deep metabolomics analysis strategy with integrated computational approaches for sorting tandem mass spectral features and annotating detected metabolites (**Figure 1**). Metabolite annotation mainly contains two complementary approaches by referring to 1) experimental reference mass spectral data collected from public databases; and 2) *in silico* reference mass spectral data generated from structural databases for biologically relevant compounds. We further characterized the structure and potential function of novel metabolites without reference in above libraries, using structural motif search combined with neutral loss scanning and metabolite association network (*Methods*).

**Figure 1.**
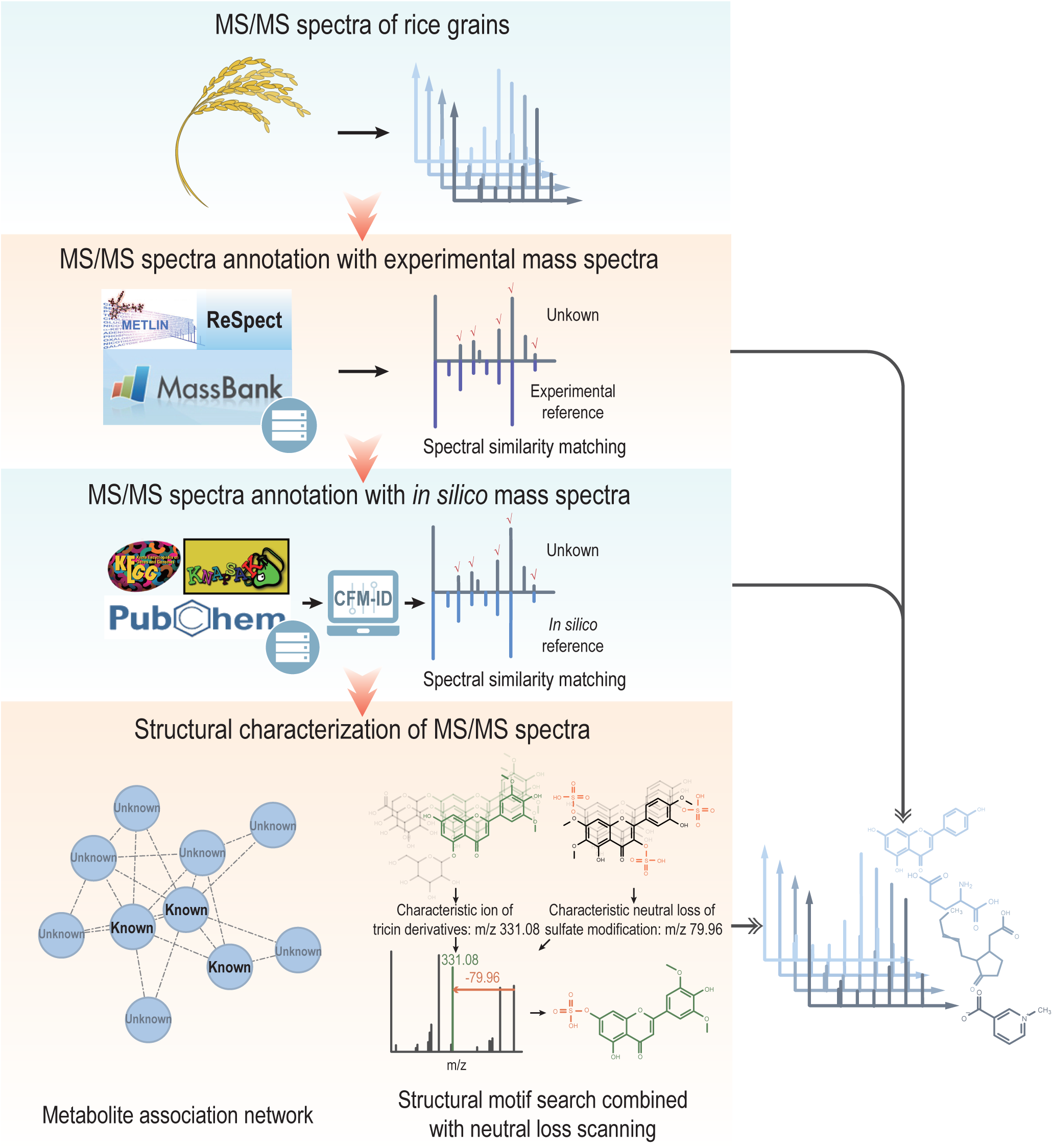
The deep metabolomics analysis strategy for large-scale structural annotation. The first approach adopted experimental reference mass spectra collected from public databases, Metlin, MassBank, and ReSpect, to annotate detected metabolites. The second approach adopted *in silico* reference mass spectra predicted from biologically relevant structure databases, KEGG, PubChem, and KNApSAcK, to annotate detected metabolites with improved coverage. CFM-ID software was used for *in silico* MS/MS spectra prediction. Two advanced methods were performed to characterize novel metabolites without reference in above spectral and structural databases. The metabolite association network was constructed to infer the structure of unknowns based on known compounds within a common cluster of related metabolites. The structural motif search combined with neutral loss scanning method was implemented to characterize the substructure of novel metabolites, by matching unknown mass spectra with characteristic fragment ion and neutral loss of specific skeletons and modifications.

The first annotation approach took advantage of the collections of experimental reference mass spectral data from public databases, including Metlin [16], MassBank [15], and ReSpect [14] (*Methods*). We evaluated the performance of two spectral similarity scoring algorithms, Normalized Dot Product (NDP) [30] and INCOS [31], and chose INCOS for subsequent analysis because of its better performance (Figure S1A). Because of the limited availability of experimental reference mass spectra, the second approach was adopted to extend the coverage with *in silico* mass spectral data for annotating those metabolites without a hit in the first approach. The *in silico* mass spectra was generated from in-house structural database (<u>S</u>tructural <u>D</u>atabase of <u>B</u>iologically <u>R</u>elevant <u>C</u>ompounds, SDBRC, Table S1) that contains the structural information of over 80,000 biologically relevant compounds collected from KEGG [32], PubChem [33], and KNApSAck [34] databases (*Methods*). The program, CFM-ID [18, 35], was used for *in silico* fragmentation of compounds from SDBRC and similarity scoring of query and reference mass spectra.

To evaluate the performance of above approaches, we sampled experimental mass spectra from Metlin and Massbank as query sets (Table S2). In first approach using experimental mass spectra as reference, INCOS had identification rates from 75 to 79% for top 1 match, and 96 to 97% when top 5 matches were included, which are comparatively higher than those of NDP (Figure S1A). In second approach using *in silico* mass spectra as reference, its performance was evaluated against KEGG and SDBRC libraries, respectively. The identification rates from 52 to 73% were observed for top 1 match, and 86 to 96% when top 5 matches were included (Figure S1B). Searching against SDBRC results in lower identification rates than KEGG. The ubiquitous isomeric compounds generally have highly similar mass spectra and are difficult to distinguish through mass spectrometry analysis. The identification rate will drop when we search against larger reference database, mainly due to more isomers contained in database [36]. SDBRC contains more biologically relevant compounds and will provide valuable reference structural information for more metabolite features. The combination of this two approaches will greatly expand the annotation coverage of plant metabolomes, and is instrumental in our study on the exploration of phytochemical diversity and function in rice.

### Constructing and annotating MS2T library for cultivated rice

To construct a MS2T library for metabolomics analysis of rice grains, we used a collection of 150 representative rice accessions (Table S3). Rice grains were harvested from farm lands in southeastern and northeastern China, and were mixed (referred as reference mixture) for subsequent processing. The extracts were subjected to UPLC-Q-Orbitrap mass spectrometer (*Methods*). The raw data from repeated analyses were aligned using Compound Discover software (v2.0, Thermo Scientific). 158,840 and 118,077 signals detected from positive and negative modes were grouped to 11,263 and 6495 merged compound features, respectively. After the quality control and redundancy filtering steps, 2637 and 2446 metabolite features were retained for positive and negative modes, respectively, in which 2234 and 2123 were tagged with MS2 spectra. Finally, these metabolite features from positive and negative modes were merged, resulting in 4491 metabolite features with 3832 tagged with MS2 spectra (Figure S2 and Tables S4, S5). These metabolite features in rice MS2T library were then annotated with our deep metabolomics analysis strategy (*Methods*). 298 metabolite features were annotated using experimental mass spectra as reference. For rest 3534 metabolite features, 1641 were annotated using *in silico* mass spectra as reference. Taken together, 1939 metabolite features were annotated in MS2T library for rice grains (Table S5). The MS2T library constructed by our study was reported as recommended [37] (Tables S4 and S5).

Benefit from the high-resolution MS and deep metabolomics analysis strategy with integrated computational approaches, we expanded the metabolite annotation coverage of rice cultivars in comparison with previous studies [2, 5, 26, 27]. Flavonoids account for a large portion of increase of annotated metabolites in rice grains. The flavonoids annotated in our study display various modifications, such as glycosylation, acetyl-glycosylation, and sulfation. The glycosylation contains monoglycoside, diglycoside, and hexuronide. Examples include RSM04010p (quercetin-3-glucoside), RSM04966p (isovitexin-7-O-xyloside), RSM05128p (apigenin-7-O-gentiobioside), RSM05322p (demethoxycentaureidin-7-O-rutinoside), and RSM02409n (apigenin 4’-glucuronide) (**Figures 2A,B,C,D,E**). The acetyl-glycosylation contains aliphatic and aromatic acylated glycoside. Examples include RSM05065p (tricin 7-(6-malonylglucoside)), RSM05648p (isovitexin 7-O-(6”’-O-E-p-coumaroyl) glucoside), and RSM05758p (7-O-(6-feruoylglucosyl) isoorientin) (**Figures 2G,H,I**). For sulfated flavonoids, an uncommon type of flavonoids, we found RSM02011n (ombuin 3-O-sulfate) (**Figure 2F**). These modifications make flavonoids diverse in solubility, reactivity, stability, and function [38, 39]. The flavonoids annotated in our study contribute to deepening our understanding of the diversity of enzymatic modifications in rice, which is benefit to the exploration of molecular mechanism of metabolite modifications in the growth, development and interaction with the environment of plants.

**Figure 2.**
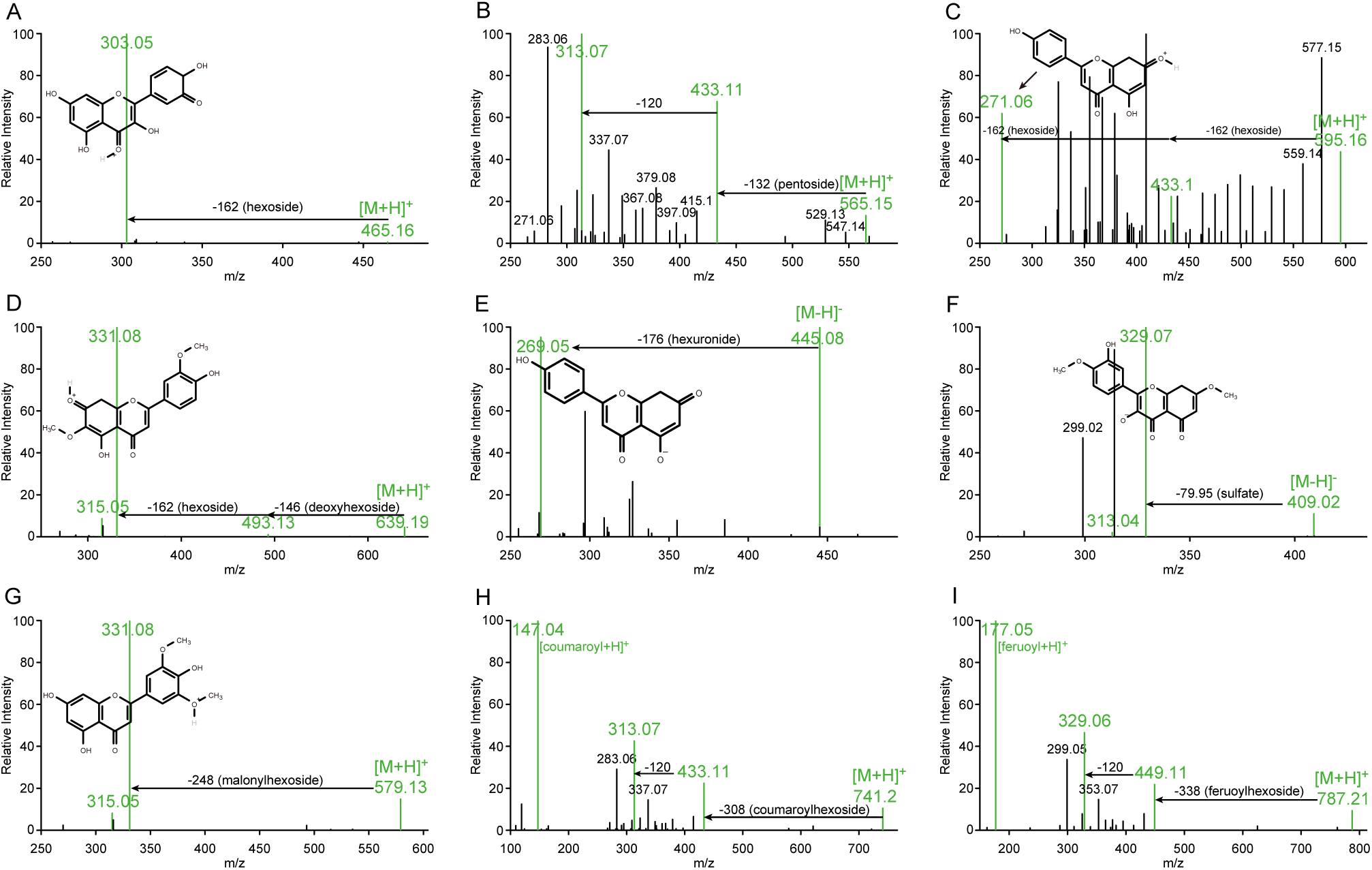
The mass spectra of annotated flavonoids with diverse modifications. [M+H]^+^ and [M-H]^-^ indicate the protonated and deprotonated precursor ion of flavonoids, respectively. RSM*****p/n indicates the serial number of rice’s screening mass spectra acquired in positive or negative ion mode. **A.** RSM04010p (isoquercitrin): the m/z 303.04922 is the featured protonated ion of quercetin, and the neutral loss of m/z 162.1087 corresponds to a hexoside group. **B.** RSM04966p (isovitexin-7-O-xyloside): the m/z 313.07016 and 433.11319 are the featured protonated ions of isovitexin (the neutral loss of m/z 120.043 is the characteristic of C-hexosyl flavonoids), and the neutral loss of m/z 132.0404 corresponds to a pentoside group. **C.** RSM05128p (apigenin-7-O-gentiobioside): the m/z 271.05936 is the featured protonated ion of apigenin, and the neutral loss of m/z 324.1032 corresponds to two hexoside groups. **D.** RSM05322p (demethoxycentaureidin-7-O-rutinoside): the m/z 315.04944 and 331.08078 are the featured protonated ions of demethoxycentaureidin, and the neutral loss of m/z 146.0587 corresponds to a deoxyhexoside (rhamnoside) group. **E.** RSM02409n (apigenin 4’-glucuronide): the m/z 269.04575 is the featured deprotonated ion of apigenin, and the neutral loss of m/z 176.0317 corresponds to a hexuronide group. **F.** RSM02011n (ombuin 3-O-sulfate): the m/z 313.03574 and 329.06674 are the featured deprotonated ions of ombuin, and the neutral loss of m/z 79.95658 corresponds to a sulfate group. **G.** RSM05065p (tricin 7-(6-malonylglucoside)): the m/z 315.04868 and 331.08017 are the featured protonated ions of tricin, and the neutral loss of m/z 248.0524 corresponds to a malonylhexoside group. **H.** RSM05648p (isovitexin 7-O-(6”’-O-E-p-coumaroyl) glucoside): the neutral loss of m/z 308.0898 corresponds to a coumaroylhexoside group, and the m/z 147.04376 is the featured protonated ion of p-coumaroyl unit. **I.** RSM05758p (7-O-(6-feruoylglucosyl) isoorientin): the m/z 449.10651 and 329.06485 are the featured protonated ions of isoorientin, the neutral loss of m/z 338.0989 corresponds to a feruoylhexoside group, and the m/z 177.05418 is the featured protonated ion of feruoyl unit.

### Differential metabolic profiles analysis revealing the featured metabolites of *indica* and *japonica* cultivars

To characterize the metabolic profiles of grains for diverse rice cultivars and understand their natural variation, we performed the untargeted metabolomics analysis on 59 rice cultivars, including 40 *indica* and 19 *japonica* (*Methods*). The metabolic profiles of rice cultivars contain the relative abundance of 3409 metabolite features (Table S6). The metabolic profiles of 59 rice cultivars were clustered based on the relative abundance of 3409 metabolite features, which displayed the differential patterns between *indica* and *japonica* cultivars (**Figure 3A**). In tree view (**Figure 3B**), the relation between *indica* with *japonica* cultivars is generally consistent to the phylogenetic relationship [40]. Through principal component analysis (PCA), *indica* and *japonica* cultivars were separated by first component (PC1) and second component (PC2), indicating the systematic difference in metabolic profiles between two subspecies (**Figure 3C**). We further performed orthogonal partial least squares discriminate analysis (OPLS-DA) to investigate the featured metabolites that differentiate *indica* and *japonica* cultivars. The *indica* and *japonica* cultivars were separated in two distinct clusters with our OPLS-DA model (**Figure 4A**). Metabolites with variable importance in projection (VIP) value greater than 2.5, were defined as featured metabolites in our study. Among 58 featured metabolites (Table S7), 11 flavonoids, 3 terpenoids, and 2 phenylpropanoids were annotated. Particularly, three novel tricin derivatives, RSM03724n (tricin-O-sulfatohexoside), RSM04661n (tricin-O-acetylrhamnoside-O-diacetylrhamnoside), and RSM05814p (tricin-O-feruloyhexoside-hexoside) (Figures S3A,B,C and Table S7), were characterized using structural motif search combined with neutral loss scanning (*Methods*).

**Figure 3.**
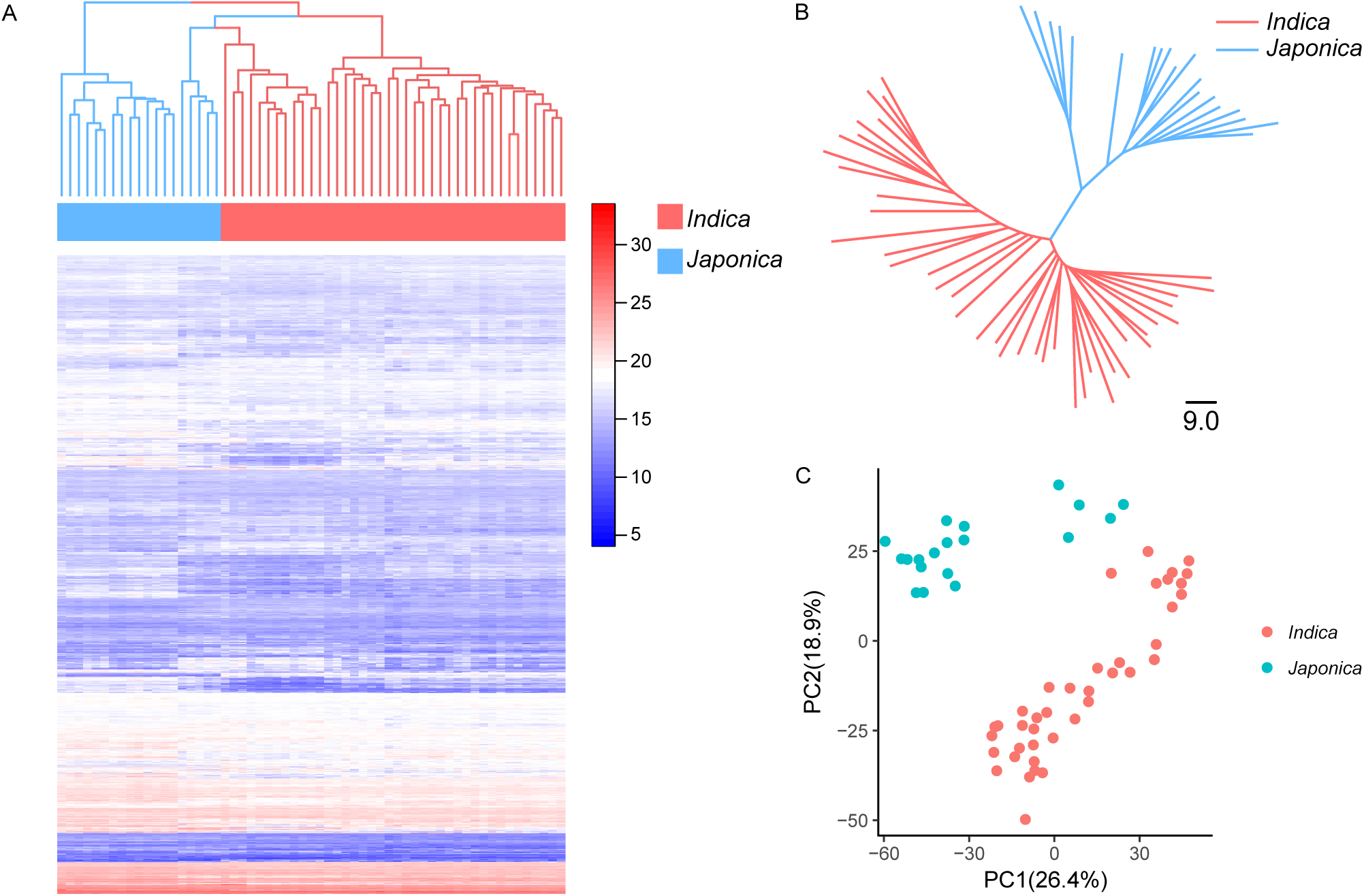
The differential metabolic profiles between *indica* and *japonica* cultivars. **A.** The heatmap and hierarchical clustering of 59 rice cultivars based on the relative abundance of 3409 metabolites. **B.** The Neighbor-joining tree of 59 rice cultivars based on the relative abundance of 3409 metabolites. **C.** The score plot for PCA of 59 rice cultivars based on the relative abundance of 3409 metabolites. The first and second principle components account for 26.4% and 18.9% of variance, respectively.

**Figure 4.**
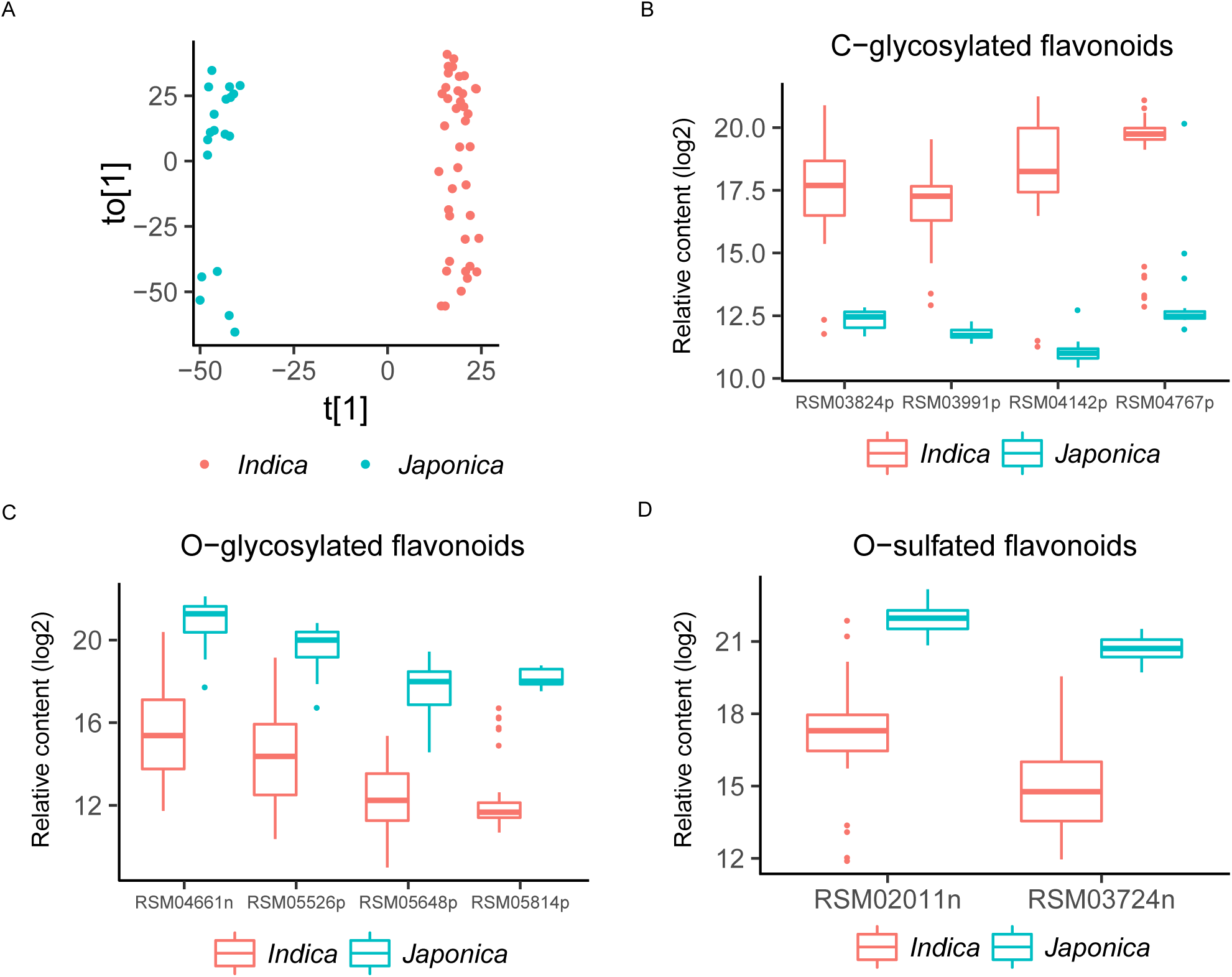
The featured metabolites of *indica* and *japonica* cultivars. **A.** The score plot for OPLS-DA of 59 rice cultivars based on the relative abundance of 3409 metabolites. The t[1] and to[1] indicate the predictive component and orthogonal component of OPLS-DA model, respectively. The R^2^X, R^2^Y (goodness-of-fit parameter), and Q^2^ (predictive ability parameter) of OPLS-DA model are 0.555, 0.99 and 0.98. **B.** The boxplot of relative abundance of four C-glycosylated flavonoids among featured metabolites. RSM03824p, cytisoside; RSM03991p, trihydroxy-methoxyflavone C-hexoside; RSM04142p, precatorin I; RSM04767p, di-C,C-pentosyl-apigenin. **C.** The boxplot of relative abundance of four O-glycosylated flavonoids among featured metabolites. RSM04661n, tricin O-acetylrhamnoside-O-diacetylrhamnoside; RSM05526p, tricin 4’-O-(guaiacylglyceryl) 7”-O-glucopyranoside; RSM05648p, isovitexin 7-O-(6”’-O-E-p-coumaroyl) glucoside; RSM05814p, tricin O-feruloylhexoside-O-hexoside. **D.** The boxplot of relative abundance of two O-sulfated flavonoids among featured metabolites. RSM02011n, ombuin 3-O-sulfate; RSM03724n, tricin O-sulfatohexoside.

We further observed the differential accumulation patterns of C-glycosylated, O-glycosylated, and O-sulfated flavonoid derivatives among featured metabolites. The levels of four C-glycosylated flavonoids (flavone C-hexoside and flavone C-pentoside), RSM03824p (cytisoside), RSM04142p (precatorin I), RSM03991p (trihydroxy-methoxyflavone C-hexoside) (Figure S3D), and RSM04767p (di-C,C-pentosyl-apigenin) (Figure S3E) are significantly higher in *indica* than *japonica* cultivars (**Figure 4B** and Table S7). In contrast, the levels of four O-glycosylated flavonoids with guaiacylglyceryl or acyl modification, RSM05526p (tricin 4’-O-(guaiacylglyceryl) ether 7”-O-glucopyranoside), RSM05648p (isovitexin 7-O-(6”’-O-E-p-coumaroyl)glucoside), RSM04661n(tricin O-acetylrhamnoside-O-diacetylrhamnoside), and RSM05814p (tricin O-feruloylhexosyl-O-hexoside) are significantly higher in *japonica* than *indica* cultivars (**Figure 4C** and Table S7). Furthermore, differences in two O-sulfated flavonoids, RSM02011n (ombuin 3-O-sulfate) and RSM03724n (tricin O-sulfatohexoside), were observed between *indica* and *japonica* cultivars (**Figure 4D** and Table S7). The differential accumulation patterns of C-glycosylated and O-glycosylated flavonoids in rice grains were consistent with previous studies in rice leaves [26, 41]. Additionally, we expanded our findings to O-sulfated flavonoids, an uncommon variety of flavonoid derivatives catalyzed by sulfotransferases [39, 42]. It has been revealed that the natural variation of salicylic acid sulfotransferase encoding gene cause the differentiation of resistance to rice stripe virus between *indica* and *japonica* subspecies [43], highlighting the significant role of sulfation in pathogen resistance of rice. However, rare study has been performed to characterize flavonoid sulfotransferases in rice. The differential accumulation patterns of O-sulfated flavonoids revealed by our study provided new insight to the natural variation of flavonoid sulfotransferase activity, which is benefit to the exploration of biosynthesis genes of flavonoid sulfotransferases and their potential functions in pathogen resistance of rice.

### Constructing metabolite association network and uncovering diverse flavonolignans from rice grains

Network-based analysis is widely used in metabolomics studies for understanding of metabolite interaction, structural characterization, and pathway elucidation [20, 44-47]. Previous studies suggested that metabolites with similar structure generally display correlation in their abundance, so the structure of unknown metabolites can be inferred by knowns through metabolite association network [2, 4, 27, 48].

We constructed the metabolite association network with Gaussian graphical model (GGM) [49], using the metabolic profiles of 59 rice cultivars (*Methods*, Figure S4A and Table S8). This network contains 2874 nodes (metabolites) with 42,147 significant edges (metabolite pairs). The 64 clusters were isolated (Table S9) from GGM network using Molecular Complex Detection (MCODE) program [50] (*Methods*). A subgroup of the first-ranked cluster mainly contains flavonoids. Besides, within the second-ranked cluster, a large number of nodes were annotated as terpenoids, most of which are triterpenoids (Figure S4B and Table S10).

A subgroup of the first-ranked cluster contains 32 metabolites (**Figure 5A** and Table S10). 13 of them were annotated as common flavonoids with hydroxy and methoxy groups (Figure S5). Notably, within this cluster, we found some flavonolignans (**Figure 5B** and Figure S6), which are produced via oxidative coupling between flavonoids with three varieties of monolignols, p-coumaryl, coniferyl, and sinapyl alcohols [51]. RSM04702p (Salcolin B) [52] and RSM04355p (5’-Methoxyhydnocarpin-D) [53] are guaiacyl flavonolignans, and RSM04382p (aegicin) is p-hydroxyphenyl flavonolignans [54]. Based on above findings, we suggested that there are other flavonolignans within this cluster. We then observed the precursor ion and fragmentation pattern of unknowns within this cluster and characterized more flavonolignans. The RSM04691p displays same fragment ion (m/z at 315.04895) with RSM04355p in mass spectra, which means they have same flavonoid moiety in structures. The mass difference between their precursor ions is 30.01031, corresponding to a methoxy group. Thus the RSM04691p has an additional methoxy group at the coniferyl alcohol moiety of RSM04355p, which was characterized as palstatin [55], a syringyl flavonolignan. With same method, the structures of RSM05474p, RSM05479p, RSM05574p, and RSM04546n were characterized. The RSM05474p was characterized as tricin O-[guaiacyl-(O-p-coumaroyl)-glyceryl] ether [56], which has an additional coumaroyl unit, a featured modification of lignins [57], at the guaiacylglyceryl group of RSM04702p. The RSM05479p, RSM05574p, and RSM04546n were characterized as tricin-oligolignols trimers, which are formed by further chain extension through oxidative coupling between tricin-oligolignols dimer (RSM04702p) and p-coumaryl alcohol or coniferyl alcohol via ether or furan bridge (Figure 5B and Figures S6E,F,G) [58]. In previous studies, the presence of unusual catechyl lignins derived from caffeyl alcohol had been revealed in plants [59]. Unexpectedly, in our study, we found that RSM04164p and RSM04201p show the spectrum features of catechyl flavonolignans, which both have the characteristic fragment ion of flavone moiety and neutral loss of caffeyl alcohol unit (m/z 166.0626). Thus we inferred that the structures of RSM04164p and RSM04201p are dihydroxy-dimethoxyflavone and tetrahydroxy-methoxyflavone moiety linked with a caffeyl alcohol unit by dioxane bridge, respectively (Figures S6I,J).

**Figure 5.**
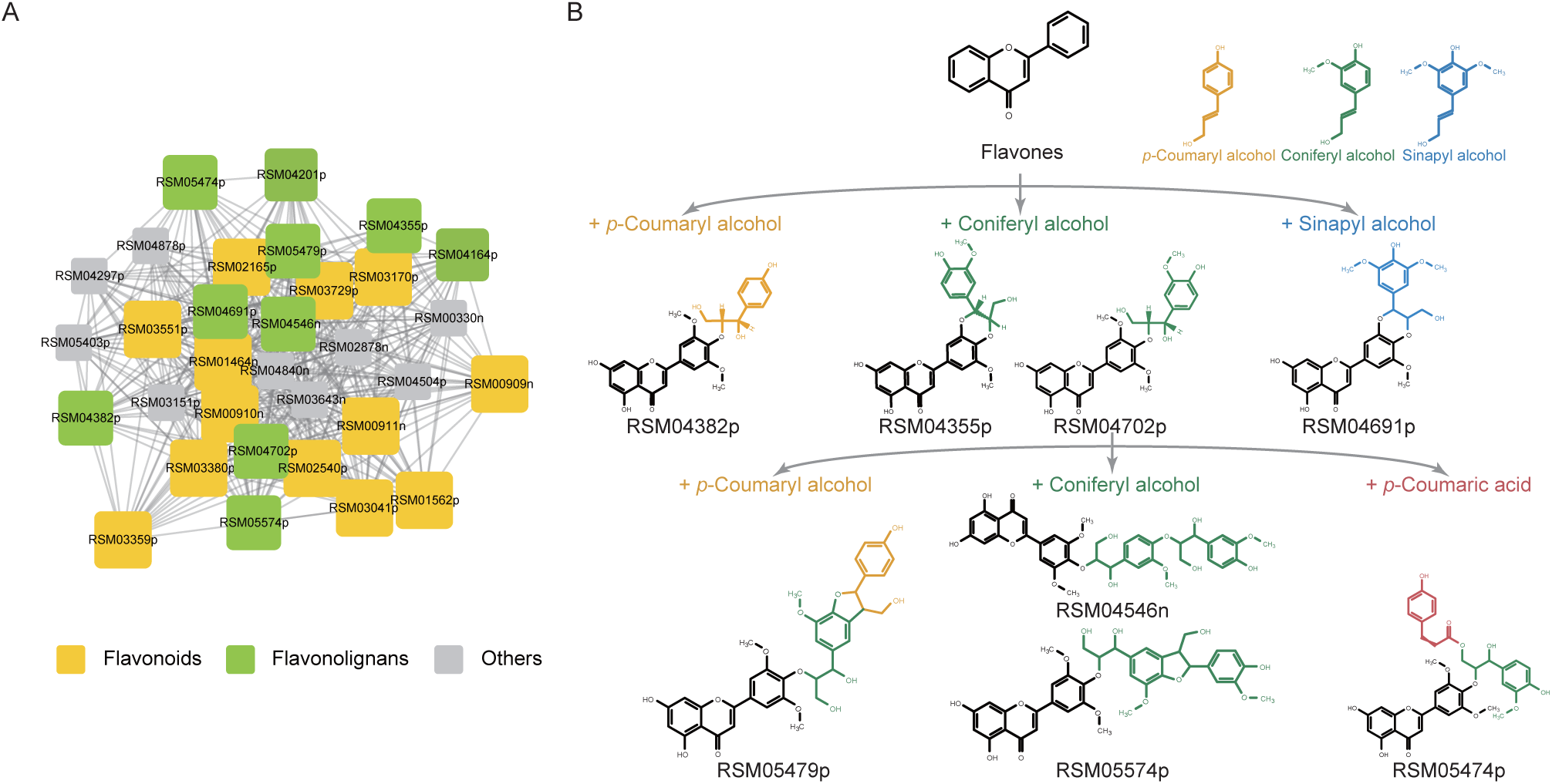
The subgroup of the first-ranked cluster containing flavonoids and flavonolignans. **A.** Components and their partial correlation relationships within the subgroup of the first-ranked cluster. **B.** The structure and relationship of characterized flavonolignans. Colors in yellow, green, and blue denote p-coumaryl alcohol, coniferyl alcohol, and sinapyl alcohol or their derived moieties, respectively. Color in red denotes the coumaroyl unit. RSM04382p, RSM04355p, RSM04702p, and RSM04691p are flavonolignans dimers derived from the oxidative coupling between flavonoids with monolignols. RSM05474p has an additional coumaroyl unit on the guaiacylglyceryl group of RSM04702p. RSM05479p, RSM05574p, and RSM04546n are flavonolignans trimers derived from the further chain extension between RSM04702p with monolignols.

In addition to RSM04382p and RSM04702p found in rice leaves and grains previously [4, 26, 60], the rest eight flavonolignans were characterized by our study in rice grains, which greatly expanded the diversity of flavonolignans in rice. Previously, the occurrence of tricin in lignins has been reported in a series of monocots [61]. Tricin was found to be incorporated into lignins as tricin-oligolignols, and acts as a nucleation site in the initiation of lignin polymers in maize [51, 58]. A group of closely-related tricin-oligolignols dimers and trimers found in our study further supported the crucial role of tricin-oligolignols in lignification of rice. Additionally, the characterization of non-tricin flavonolignans, such as RSM04355p and RSM04691p, provided evidence for the presence of more diverse flavonoids in lignification. Within this cluster, six additional metabolites were found to contain featured ions of tricin in their mass spectra, which may be putative tricin derivatives. Two of them show the neutral loss of guaiacylglyceryl or p-hydroxyphenylglyceryl unit, although their entire structures remain unknown (Figure S7 and Table S10).

## Conclusions

The technical and analytical obstacles in the identification of metabolites hindered the further research of phytochemical diversity and function in plants. To address the issue of low identification coverage in plant metabolomics, we adopted a deep metabolomics analysis strategy for large-scale metabolite structural annotation. The experimental and *in silico* mass spectra was used to facilitate the metabolite annotation with high coverage. The structural motif search combined with neutral loss scanning, and metabolite association network methods were further adopted to characterize the structure and function of metabolites in rice. The untargeted metabolomics study on rice grains was performed, and the coverage of annotated metabolites was significantly improved. Benefited from the rice metabolome with expanded annotation coverage, the systematic differences in metabolic profiles between *indica* and *japonica* cultivars were further defined, including the differential accumulation patterns of C-glycosylated, O-glycosylated, and O-sulfated flavonoids, and a series of closely-related flavonolignans with key roles in lignification were uncovered. Our strategy can be applied to the metabolomics researches of other agronomically important plants, with great potential in the enhancement of crops quality and nutrition value through genetic breeding.

## Materials and methods

### Plant materials

Rice cultivars, including 93 *japonica* and 85 *indica* accessions, were used in this study (Table S3). Rice cultivars were planted and harvested during the summer season in 2015 and 2016 from two locations of China: farm lands in Jiangsu (Yangzhou, E 119°53’, N 32°42’, southeastern China) and in Heilongjiang (Harbin, E 126°C 53’, N 45°69’, northeastern China). Rice cultivars were planted in the field, ten plants for each row, and three rows for each accession.

For each accession, grains were harvested for two biological replicates, each containing grains from three individual plants. Grains were packed in gauze bag and air-dried in shade. Two grams of dried grains were ground using tissue grinder (catalog No. 05010997; Shanghai BiHeng Biotechnology Company Limited; Shanghai; China) at 55Hz for 40 seconds. The fine powder for each accession was stored at -80°C for subsequent processing.

### Chemicals

HPLC grade methanol, acetonitrile and acetic acid were obtained from Merck Company (catalog No. 1.06007.4008, 1.00030.4008, 5.43808.0250; Merck KGaA; Darmstadt; Germany). Ultra-pure water was produced using Millipore water purifier (Milli-Q; Millipore; Billerica; MA; USA). The lidocaine and lincomycin (CAS No. 137-58-6, 859-18-7; Dr. Ehrenstorfer, GmbH; Augsburg; Germany) were purchased from ANPEL Laboratory Technologies (Shanghai) Inc.. Other chemicals were purchased from Sigma-Aldrich (Shanghai) Trading Co., Ltd. (Sigma-Aldrich; Merck KGaA; Darmstadt; Germany), if not otherwise specified.

### Metabolite extraction

150 mg of powder of rice grains was mixed with 1.5 mL 70% aqueous methanol solution A (containing 1 mg/L vitexin, 1 mg/L p-coumaric acid, and 1 mg/L lidocaine as internal standards). The mixture was vortexed every 10 min for 3 times and placed in 4°C refrigerator overnight. The mixture was then centrifuged at 12,000g for 10 min in 4°C. The supernatant of mixture was dried with concentrator under vacuum and re-dissolved with 150 uL 70% aqueous methanol solution B (containing 1 mg/L capsaicin and 1 mg/L lincomycin as internal standards). Then the extract was filtered with 0.22μm filter (catalog No. SCAA-104; ANPEL Laboratory technologies Inc; Shanghai; China) and transferred into sample bottle for the subsequent UPLC-MS/MS analysis.

### UPLC-MS/MS analysis

Chromatographic separation of extract samples was performed on Waters Acquity Ultra Performance Liquid Chromatography using an ACQUITY UPLC BEH C18 column (pore size: 1.7μm, length: 2.1*100mm) (Waters Corporation; Milford; MA; USA). The mobile phase consisted of (A) water with 0.04% acetic acid and (B) acetonitrile with 0.04% acetic acid. The gradient program was as follows: 95:5 A/B at 0 min, 5:95 A/B at 20.0 min, 5:95 A/B at 24.0 min, 95:5 A/B at 24.1 min, and 95:5 A/B at 30.0 min. The flow rate was 0.25 mL/min and the injection volume was 5 μL. The column temperature was 40°C.

The UPLC system was coupled with Q Exactive™ Hybrid Quadrupole-Orbitrap High Resolution Mass Spectrometer (Q-Orbitrap-HRMS) (Thermo Fisher Scientific; Waltham; MA; USA). The MS acquisition was performed in positive and negative ionization with FullScan/dd-MS2 (top 8) mode, in which the MS/MS spectra of most abundant ions (top 8) within each scanning window was automatically obtained. MS full scan mass resolution was set to 70,000 at m/z 200 and data-dependent MS/MS with full scan mass resolution was reduced to 17,500 at m/z 200. The m/z range of MS full scan was 100-1000.

Heated electrospray ionization (HESI) parameters were as follows: Spray voltage (+), 4000 V; Spray voltage (-), 3500 V; Capillary temperature, 320°C; Sheath gas, 35 arb; Aux gas, 8 arb; Probe heater temperature, 350°C; S-Lens RF level, 50. Higher energy Collisional Dissociation (HCD) energies were 15eV and 40eV, and average MS/MS spectrum was obtained. The mass spectrometer was calibrated using Pierce™ LTQ Velos ESI positive ion Calibration Solution and Pierce™ ESI negative ion Calibration Solution (Thermo Fisher Scientific; Waltham; MA; USA).

The sequence of injections for extract samples was randomized to reduce bias. The grains mixture of 150 randomly selected rice accessions was used to build a reference MS2T library. Reference mixture was submitted to UPLC-MS/MS system once every 10 samples. In total, injections of reference mixture were repeated for 43 times in positive and negative modes.

### Mass spectrum data processing

The raw data generated from HESI-Q-Orbitrap-HRMS were processed with Compound Discoverer software (version 2.0, Thermo Scientific) using its automatic workflow. The retention time aligning parameters were as follows: mass tolerance, 5ppm; maximum Shift, 0.5min. The unknown compounds detecting parameters were as follows: min peak intensity, 2E6; S/N threshold, 5.

Raw metabolite features were further filtered by: 1) removing signals that are of poor quality or non-biological origin [62], i.e. features with reproducibility <90%, sample to blank ratio <10%, relative standard deviation (RSD) >50%, or peak area less than 1E5; 2) removing redundancy from multi-ion adducts (Na^+^, K^+^, NH4^+^, Cl^-^), isotopes, in-source fragmentation, or dimerization. Metabolite features in positive ([M+H]^+^) and negative ([M-H]^-^) modes were merged with following parameters: exact mass tolerance, 5ppm; and retention time tolerance, 0.5min. The in-house script based on Xcalibur Development Kit (XDK) in Xcalibur software (version 2.2, Thermo Scientific) was used to automatically extract MS2 spectra of metabolite features.

### Metabolite annotation

Metabolite annotation mainly adopted two complementary approaches with experimental/*in-silico* mass spectra as reference. The first approach used the experimental reference mass spectra library collected from public databases, such as Metlin [16], MassBank [15], and ReSpect [14]. This library contained a total of 98,658 mass spectra for about 24,385 compounds. Two algorithms, NDP and INCOS were implemented as described [30, 31] using perl scripts to score the similarity between query and reference mass spectra. INCOS algorithm was selected for further analysis because of its better performance. We respectively searched against the Metlin, MassBank, and ReSpect libraries and merged the annotation results subsequently. The experimental reference mass spectra that have similar precursor m/z (mass tolerance: 10 ppm) with query mass spectra were retrieved, and were compared for similarity using INCOS. Reference mass spectra with similarity score >0.75 was retained for the annotation of query mass spectra. The reference spectra with the highest similarity score in the annotation results was selected as the putative annotation for the query spectra. In the evaluation of NDP and INCOS algorithms using query spectra sampled from Metlin or MassBank, the query spectra themselves were excluded from the matching results to rule out bias. The performance of the first annotation approach (INCOS) with similarity score cutoff (0.75) was further evaluated with the test set for standard MS/MS spectra of Fiehn HILIC Library from MassBank of North America (Figure S8A and Table S11).

The second approach was adopted to extend the annotation coverage with *in silico* mass spectra. First, the structure data were collected from three biologically relevant structure databases, including KEGG, ‘BioChem’ (the manual selected subset of biologically relevant compounds in PubChem), and KNApSAck. For ‘BioChem’ database, compounds in PubChem with NCBI BioSystems annotation, biological role classification of ChEBI Ontology, Flavonoids or Prenol Lipids classification of LIPID MAPS [63] were selected. The OpenBabel software [64] was used to convert the raw structural data to machine-readable structural information, including formula, exact mass, Simplified Molecular-Input Line-Entry System (SMILES), and the IUPAC International Chemical Identifier (InChI). Through merging compounds from different databases and removing redundancy, we constructed a structural database that contains 85,342 non-redundant compounds (SDBRC), which was used as reference to retrieve and generate *in silico* mass spectra. The program, CFM-ID, was used for *in silico* fragmentation of compounds from SDBRC, and similarity scoring as described [18, 35]. CFM-ID adopts machine learning technique with probabilistic generative model for compound fragmentation process. The source code for CFM-ID software (version 2.0) was obtained from SourceForge platform (https://sourceforge.net/projects/cfm-id/), and compiled on Linux system (CentOS release 6.2). The *in silico* mass spectra for reference compounds that have similar mass (mass tolerance: 5 ppm) with query mass spectra were generated using CFM-ID, and similarity scores between query and *in silico* mass spectra were calculated. Reference compounds with similarity score >0.3 were retained for the annotation of query mass spectra. The performance of the annotation through CFM-ID with similarity score cutoff (0.3) was further evaluated with the test set for standard MS/MS spectra of Fiehn HILIC Library from MassBank of North America (Figure S8B and Table S11).

The annotation results for 17 metabolite features were further identified through the comparison of retention time (RT) and MS/MS spectra with standard compounds (Figure S9 and Table S12).

### Structural motif search combined with neutral loss scanning

The structural motif search combined with neutral loss scanning, was further developed from previous studies [14, 26, 60, 65]. It is based on the theory that compounds with similar structures (i.e. same skeletons or modifications) would generate featured fragment ions or neutral losses in mass spectral analysis. These compounds often belong to a certain phytochemical class. Flavonoids have a core diphenylpropane backbone (C6-C3-C6) with diverse modifications and display regular fragmentation patterns in their mass spectra. In order to mine their fragmentation regularities systematically and facilitate the characterization of novel flavonoids, 3145 MS/MS spectra of two major classes of flavonoids, flavones and flavonols, were generated *in silico* by CFM-ID software from structure data in LIPID MAPS Structure Database [63]. Through statistical analysis, we obtained a series of structural motifs (characteristic fragment ions) frequently found in mass spectra of flavones and flavonols, such as m/z at 287.0550145 (featured ion of kaempferol derivatives), m/z at 303.0499291 (featured ion of quercetin derivatives), m/z at 271.0600999 (featured ion of apigenin derivatives), and m/z at 301.0706646 (featured ion of chrysoeriol derivatives). In addition, a set of frequently found neutral losses were observed, such as the neutral losses of hexoside (m/z 162.0530308), pentoside (m/z 132.0423309), rhamnoside (m/z 146.0576808), hexuronide (m/z 176.0322455), sulfate (m/z 79.9568149), and coumaroylhexoside (m/z 308.0892455) groups. We searched for the presence of structural motifs and neutral losses in unknown MS/MS spectra to characterize its putative structure. The detailed steps of structural motif search combined with neutral loss scanning were listed in Figure S10.

### The metabolic profiles of rice cultivars

The metabolic profiles of 59 rice cultivars grown in Yangzhou in 2016 was obtained from the corresponding peak areas of raw mass spectrometric data using Compound Discoverer software (v2.0, Thermo Scientific). The metabolic profiles of rice cultivars were defined according to our reference MS2T library. The metabolite features in metabolic profiles were aligned with reference MS2T library to determine corresponding structural information as described [65]. The parameters used to determine corresponding structural information were as follows: the tolerance of retention time, 0.35min; and the tolerance of mass, 5ppm. To ensure the consistency among samples during UPLC-MS/MS analysis, the reference control mixtures were inserted into the analytical sequence once every 10 samples. The data of metabolite abundance was normalized based on internal standard and reference control mixtures as described [66]. Two biological repeats for each rice accession were performed, and the normalized data was averaged and log2-transformed for further analysis. The detailed steps of the acquisition and processing of relative abundance data of metabolites were listed in Appendix S1.

### Construction of GGM-based network

For construction of GGM network, a data matrix containing the relative abundance of 3409 metabolite features for 59 rice cultivars was first generated. GeneNet package [67] was used to calculate the partial correlation coefficients and test the significance of partial correlation of each metabolite pair. The metabolite pairs with probability greater than 0.99 (local fdr < 0.01) were defined as significant edges and included in GGM network. The Cytoscape software [68] was used for the visualization of GGM network. The MCODE application was used to find clusters from GGM network with parameters as defaulted [50].

### Statistical analysis

R software (version 3.2.3; https://www.R-project.org/) [69] was mainly used for statistical analysis, if not specifically indicated otherwise. The metabolic profiles of rice cultivars were clustered using hierarchical clustering. The method of hierarchical clustering is Unweighted Pair Group Method with Arithmetic mean (UPGMA). The heatmap was constructed by heatmap.2 function in gplots package (https://CRAN.R-project.org/package=gplots)[70]. The relatedness distance between metabolic profiles of rice cultivars was calculated by Euclidean distance function. The Neighbor-joining tree was constructed by MEGA7 software [71] using the matrix of Euclidean distance between metabolic profiles of rice cultivars. PCA and OPLS-DA were carried out by SIMCA-P software (version 14.0; Umetrics, Sweden).

## Data availability

The data supporting the findings of this study are available from the supplementary materials.

## Authors’ contributions

X (Xuan) L conceived and designed the project. X (Xuetong) L, HZ, NX,YS, LC, CW and AL prepared samples and performed LC-MS analysis experiments. X (Xuetong) L, HZ, XW, and YS performed bioinformatics analyses. JH and ZW advised on rice metabolomics experiment and data analysis. X (Xuan) L and X (Xuetong) L wrote and edited the manuscript. All authors approved the manuscript.

## Acknowledgements

We thank Prof. Jie Luo for assistance in sample preparation in LC-MS analysis, and Ms. Ping Chen for help with data submission. This work was supported by grants from the Special Fund for Strategic Pilot Technology of Chinese Academy of Sciences [grant number XDA24010403], the National Key Research and Development Program of China [grant number 2018YFA0900700], the Strategic Project for Biological Resources and Service Network of Chinese Academy of Sciences [grant number ZSYS-014], the Major Project of Jiangsu Province of China for Significant New Varieties Development [grant number PZCZ201702], the National Natural Science Foundation of China [grant numbers 31900470, 31701137, 31771412].

## Supplementary material

**Figure S1** The performance evaluation of two annotation approaches

**A.** Evaluating the performance of annotation with experimental mass spectra as reference. Query mass spectra were sampled from Metlin or Massbank, and spectral similarity was scored with NDP or INCOS algorithm. **B.** Evaluating the performance of annotation with *in silico* mass spectra as reference. Query mass spectra were sampled from Metlin or Massbank, and spectral similarity was scored by CFM-ID software.

**Figure S2** The workflow for MS2T library construction

Sample preparation, metabolite extraction, UPLC-HRMS analysis, and mass spectral data processing were as described (*Methods*).

**Figure S3** The mass spectra of several featured metabolites of *indica* and *japonica* cultivars

[M+H]^+^ and [M-H]^-^ indicate the protonated and deprotonated precursor ion of metabolites, respectively.

**A.** The mass spectra of RSM03724n (tricin O-sulfatohexoside). The m/z 313.03586 and 329.06653 are the featured deprotonated ions of tricin, and the neutral losses of m/z 79.95697 and 162.0536 correspond to the sulfate and hexoside groups, respectively. **B.** The mass spectra of RSM04661n (tricin O-acetylrhamnoside-O-diacetylrhamnoside). The neutral loss of m/z 418.1954 corresponds to the acetylrhamnoside and diacetylrhamnoside groups. **C.** The mass spectra of RSM05814p (tricin O-feruloylhexoside-O-hexoside). The m/z 315.04944 and 331.08084 are the featured protonated ions of tricin, and the neutral loss of m/z 500.1448 corresponds to the feruloylhexoside and hexoside groups. The m/z 177.05428 is the featured protonated ion of feruloyl unit. **D.** The mass spectra of RSM03991p (trihydroxy-methoxyflavone C-hexoside). The neutral loss of 120.0449 is the characteristic of C-hexosylflavones. **E.** The mass spectra of RSM04767p (di-C,C-pentosyl-apigenin). The neutral loss of 90.05359 is the characteristic of C-pentosylflavones.

**Figure S4** The diagram of metabolite association network

**A.** The metabolites association network of rice grains according to the metabolic profile of 59 rice cultivars. **B.** The second ranked cluster that mainly consist of terpenoids.

**Figure S5** The structure of flavonoids within the subgroup of first-ranked cluster These flavonoids have diverse numbers of hydroxyl and methoxyl groups in their structures.

**Figure S6** The structure and mass spectra of characterized flavonolignans

[M+H]^+^ and [M-H]^-^ indicate the protonated and deprotonated precursor ion of flavonolignans, respectively.

**A.** RSM04382p, aegicin, which has a structure of tricin moiety with a p-coumaryl alcohol linked by an ether bond. The m/z 315.04950 and 331.08084 are featured protonated ions of tricin, and the neutral loss of m/z 166.0628 corresponds to the p-hydroxyphenylglyceryl unit. **B.** RSM04355p, 5’-Methoxyhydnocarpin-D, which has a structure of methoxyluteolin moiety with a coniferyl alcohol linked by a dioxane bridge. The m/z 315.04874 is featured protonated ion of methoxyluteolin moiety, and the neutral loss of m/z 180.0789 corresponds to the coniferyl alcohol unit. **C.** RSM04702p, salcolin B, which has a structure of tricin moiety with a coniferyl alcohol linked by an ether bond. The neutral loss of m/z 196.0736 corresponds to the guaiacylglyceryl unit. **D.** RSM04691p, palstatin, which has a structure of methoxyluteolin moiety with a sinapyl alcohol linked by a dioxane bridge. The neutral loss of m/z 210.0891 corresponds to the sinapyl alcohol unit. **E.** RSM05479p, which has a structure of salcolin B moiety with a p-coumaryl alcohol linked by a furan bridge (characterized through its mass spectra). **F.** RSM04546n, which has a structure of salcolin B moiety with a coniferyl alcohol linked by an ether bond (characterized through its mass spectra). **G.** RSM05574p, which has a structure of salcolin B moiety with a coniferyl alcohol linked by a furan bridge (characterized through its mass spectra). **H.** RSM05474p, tricin O-[guaiacyl-(O-p-coumaroyl)-glyceryl] ether, which has an additional coumaroyl unit at the guaiacylglyceryl group of salcolin B. The neutral loss of m/z 342.1071 corresponds to the O-p-coumaroyl-guaiacylglyceryl group, and the m/z 147.04379 is featured protonated ion of p-coumaroyl unit. **I.** RSM04164p, putative catechyl-type flavonolignans characterized through its mass spectra, which has a structure of dihydroxy-dimethoxyflavone moiety with a caffeyl alcohol linked by a dioxane bridge. The m/z 313.07019 is featured protonated ion of dihydroxy-dimethoxyflavone moiety, and the neutral loss of m/z 166.0626 corresponds to the caffeyl alcohol unit. **J.** RSM04201p, putative catechyl-type flavonolignans characterized through its mass spectra, which has a structure of tetrahydroxy-methoxyflavone moiety with a caffeyl alcohol linked by a dioxane bridge. The m/z 315.04919 is featured protonated ion of tetrahydroxy-methoxyflavone moiety.

**Figure S7** The mass spectra of 6 putative tricin derivatives

[M+H]^+^ and [M-H]^-^ indicate the protonated and deprotonated precursor ion of putative tricin derivatives, respectively (see Table S10 for details).

**Figure S8** The performance evaluation of the annotation approaches with test set of Fiehn HILIC Library

The test set used for performance evaluation was collected from the MassBank of North America (see Table S11 for details). **A.** The percentages of top 3 correct annotation of the first annotation approach (INCOS). The evaluations were performed with Metlin and MassBank as reference, and the cutoff of similarity score was 0.75. **B.** The percentages of top 3 correct annotation of the second annotation approach. The evaluations were performed with the *in silico* mass spectra generated from KEGG and SDBRC database as reference, and the cutoff of similarity score was 0.3.

**Figure S9** The matching results of MS/MS spectra between 17 metabolite features with standard compounds

The MS/MS spectra for standard compounds were acquired by the same metabolic analysis methods with metabolite features in MS2T library. The detailed information for identification of 17 metabolite features were listed in Table S12.

**Figure S10** The detailed steps of structural motif search combined with neutral loss scanning

1) The raw structure data of flavones and flavonols were collected from the LIPID MAPS Structure Database. The OpenBabel software was used to convert the raw structure data into the machine-readable structural information, including formula, exact mass, Simplified Molecular-Input Line-Entry System (SMILES), and the IUPAC International Chemical Identifier (InChI). The non-redundant structure data for 3145 flavones and flavonols were obtained though merging the compounds with identical structure.

2) The theoretical MS/MS spectra for flavones and flavonols were predicted by CFM-ID software with structure data as input.

3) The fragments with top 25% of intensity in each mass spectra were retained for further analysis. The fragments with lower structure similarity (calculated by OpenBabel software) with the diphenylpropane backbone (C6-C3-C6) of flavonoids, were filtered out as non-aglycone derived fragments.

4) The remaining fragments with identical mass were merged and sorted by their frequency. The merged fragments were considered as the characteristic fragments of flavones and flavonols. The fragments with comparatively high frequency represent the structural motifs frequently found in flavones and flavonols, such as m/z at 287.0550145 (featured ion of kaempferol derivatives), m/z at 303.0499291 (featured ion of quercetin derivatives), m/z at 271.0600999 (featured ion of apigenin derivatives), and m/z at 301.0706646 (featured ion of chrysoeriol derivatives), *etc*.

5) The mass difference between precursor ion and characteristic fragments in each MS/MS spectra were calculated as neutral losses. The neutral losses with similar mass (tolerance: 10ppm) were merged and sorted by their frequency. The merged neutral losses were considered as the featured neutral losses of flavones and flavonols. The neutral losses with comparatively high frequency represent the modifications frequently occurred in flavones and flavonols, such as the neutral losses of hexoside (m/z 162.0530308), pentoside (m/z 132.0423309), rhamnoside (m/z 146.0576808), hexuronide (m/z 176.0322455), sulfate (m/z 79.9568149), and coumaroylhexoside (m/z 308.0892455) groups, *etc*.

6) The characteristic fragments and featured neutral losses were used as the queries to search unknown MS/MS spectra for structural characterization. We firstly searched for the presence of the characteristic fragments (mass tolerance: 10ppm) in unknown MS/MS spectra. And if there were characteristic fragments, the mass difference between precursor ion and each characteristic fragment was calculated. If the mass difference was similar (mass tolerance: 10ppm) with featured neutral losses, the unknown MS/MS spectra was putatively characterized as flavones or flavonols.

7) The putatively characterized structures were further inspected with other available phytochemical information to improve the confidence of structural characterization.

**Table S1** Structural Database of Biologically Relevant Compounds (SDBRC)

**Table S2** The information of test sets

**Table S3** The information of rice varieties

**Table S4** Metabolite Reporting Checklist

**Table S5** The MS2T library of rice grains

**Table S6** The relative abundance data of 3409 metabolites for 59 rice cultivars

**Table S7** The information of featured metabolites of indica and japonica cultivars

**Table S8** The Gaussian graphical model (GGM)-based network

**Table S9** The 64 clusters within GGM network

**Table S10** The information of metabolites within the clusters of GGM network **Table S11** The detailed information of the test dataset of Fiehn HILIC Library **Table S12** The detailed information of the identification with standard compounds for 17 metabolite features

**Table S13** The mass spectra of metabolites mentioned in paper

**Appendix S1** The detailed steps for the acquisition and processing of relative abundance data of metabolites

